# VASP Promotes Aortic Valve Calcification by Interacting with FBLIM1 to Activate the Focal Adhesion Pathway

**DOI:** 10.64898/2026.07.17.739282

**Authors:** Xiaolin Liu, Qingyuan Xu, Kai Xing, Nan Zhang, Qiang Zheng, Peilei Sun, Rui Li, Wenlong Zhang, Zhaoshui Li, Zhengjun Wang

**Affiliations:** Department of Cardiovascular Surgery, Shandong Provincial Hospital Affiliated to Shandong First Medical University, Jinan 250021, Shandong, China; Medical Science and Technology Innovation Center, Shandong First Medical University, Jinan 250117, Shandong, China; Department of Ultrasound, Shandong Provincial Hospital Affiliated to Shandong First Medical University, Jinan 250021, Shandong, China; Department of Cardiovascular Surgery, The First Affiliated Hospital of Shandong First Medical University & Shandong Provincial Qianfoshan Hospital, Jinan 250014, Shandong, China

**Keywords:** Calcific aortic valve disease, Vasodilator-stimulated phosphoprotein, Focal adhesions, Filamin binding LIM protein 1, Extracellular matrix

## Abstract

**Background:** Calcific aortic valve disease (CAVD) is the most prevalent valvular heart disease, yet there is no effective pharmacological therapy to halt or reverse its progression. Focal adhesions (FAs) are dynamic structures that connect cells to the extracellular matrix (ECM), serving not only as mechanical anchors but also as critical signalling hubs that regulate cell adhesion, spreading, migration, differentiation, and mechanotransduction. However, their specific role in CAVD pathogenesis remains largely unknown.

**Methods:** To identify upregulated hub genes, an integrated analysis of proteomics and RNA sequencing was performed on calcified aortic valves. To investigate functional roles in vitro, we utilized genetic knockdown and overexpression of VASP in valvular interstitial cells (VICs), followed by phenotypic evaluations of osteogenic differentiation, calcification, and elastin (ELN) secretion. Downstream mechanisms were explored using RNA-seq in VASP-overexpressing VICs, alongside pharmacological inhibition of the FA pathway. Finally, protein-protein interaction assays were conducted to map and analyze the physical binding between VASP and the structural domains of FBLIM1.

**Results:** Integrated omics analysis identified VASP as a significantly upregulated gene in human calcified aortic valve tissues. Functionally, VASP knockdown inhibited, whereas its overexpression promoted, the osteogenic differentiation of VICs. RNA-seq revealed that VASP overexpression activated the FA pathway, and the pharmacological blockade of this pathway successfully suppressed calcification both in vitro and vivo. Mechanistically, we demonstrated a direct physical interaction between VASP and the third LIM zinc-binding domain of FBLIM1, and knockdown of FBLIM1 effectively reversed the pro-calcific effects of VASP. Finally, VASP overexpression was found to promote the secretion of ELN, which subsequently contributed to the calcification process.

**Conclusion:** Our study reveals that the interaction between VASP and FBLIM1 drives CAVD progression by activating the FA pathway, which subsequently leads to excessive ELN secretion. These findings reveal a novel mechanistic pathway and may provide a potential therapeutic target for CAVD intervention.

**What are the Clinical Implications?:** We have identified the VASP-FBLIM1-FA-ELN axis as a novel molecular pathway involved in CAVD pathogenesis, greatly enriching our understanding of the complex process of aortic valve calcification. This pathway intricately links cytoskeletal dynamics, cell adhesion signalling, and ECM remodelling, providing a new theoretical framework for pathophysiological research in CAVD. In clinical practice, VASP and its downstream pathway components, particularly FBLIM1 and the FA pathway, may serve as potential biomarkers and therapeutic targets for the early diagnosis and treatment of CAVD. For example, the FA pathway inhibitors PF-573228 and Y15 demonstrated significant anti-calcification effects both in vitro and in vivo, suggesting that targeting this pathway may offer a new nonsurgical intervention strategy for patients with CAVD.

## 1. Introduction

Calcific aortic valve disease (CAVD) is a degenerative cardiac condition characterized by progressive calcium deposition, fibrotic remodelling, and functional impairment of the aortic valve and predominantly affects the elderly population^1,2^. Global epidemiological studies have reported that CAVD affects approximately 12.6 million individuals and contributes to 102,700 deaths annually^3^. Currently, clinical treatment relies heavily on invasive valve replacement surgery. Although the increased adoption of transcatheter aortic valve replacement (TAVR) has reduced mortality in high-risk patients, complications such as subclinical thrombosis and conduction abnormalities remain significant^4^. Osteogenic differentiation of valvular interstitial cells (VICs) is a key event in CAVD pathogenesis^5^. This process is intricately regulated at multiple levels, involving key signalling pathways^6,7^, epigenetic modifications (e.g., m6A methylation and lactylation)^8–11^, and metabolic dysregulation (e.g., iron overload)^12,13^. Owing to its extreme complexity, the underlying mechanism has not been fully elucidated. Therefore, an in-depth understanding of the core molecular mechanisms driving the osteogenic differentiation of VICs and the identification of intervenable key targets for early intervention and drug development in CAVD are highly important.

Vasodilator-stimulated phosphoprotein (VASP) is a crucial actin-binding protein that links various kinase signalling pathways to actin assembly and serves as a key regulator of the cytoskeleton^14^. Studies have reported that VASP expression is upregulated in neonatal and hypertrophic heart tissues in response to increased mechanical load^15^. Given that hypercholesterolemia is an independent risk factor for CAVD, it is notable that VASP expression is also increased in patients with hypercholesterolemia^16^. VASP is also a core component of focal adhesions (FAs) and plays important roles in regulating actin polymerization and cell migration^17,18^. However, its specific role in CAVD has not been clearly defined.

Focal adhesions (FAs) are dynamic structures that connect cells to the extracellular matrix (ECM)^19,20^. They serve not only as mechanical anchors but also as critical signalling hubs that regulate cell adhesion, spreading, migration, differentiation, and mechanotransduction^21–23^. VASP, as a constituent of FAs, directly participates in regulating the assembly and function of these structures^24–27^. In this study, we found that VASP is highly expressed in calcified aortic valves. It interacts with FBLIM1 to activate the FA pathway, promotes the secretion of ELN, and consequently regulates the osteogenic differentiation of VICs. Our study reveals the molecular mechanism through which the VASP-FBLIM1-FA pathway promotes CAVD progression, providing a new potential therapeutic target for the prevention and treatment of CAVD.

## 2. Methods

### 2.1 Human samples and ethics

Normal aortic valve leaflets were obtained from the excised valve leaflets of heart transplant patients or patients who underwent type A aortic dissection, whereas calcified aortic valve leaflets were collected from patients with aortic stenosis who underwent surgical aortic valve replacement. Patients with a bicuspid aortic valve, rheumatic aortic stenosis, concomitant coronary artery disease, diabetes mellitus, renal insufficiency, or infective endocarditis were excluded. All patients provided informed written consent for the use of their valve tissues in this study. The study was conducted in accordance with the Declaration of Helsinki, and the protocol was approved by the Medical Ethics Committee of Shandong First Medical University Affiliated Provincial Hospital (SZRJJ No. 2025-346). The baseline characteristics of the patients are shown in Supplemental Table S1.

### 2.2 Isolation and Culture of Primary Cells

Fresh aortic valve specimens were washed three times with PBS. The tissues were then transferred to culture dishes and digested with 2 mg/mL collagenase I (Biosharp, China) for 12 hours. Cells were collected by centrifugation. Primary human valvular interstitial cells (hVICs) were cultured in DMEM supplemented with 10% foetal bovine serum (LONSERA, Uruguay) and 1% penicillin‒streptomycin (Sangon Biotech, China). Cells were passaged when they reached 85–90% confluence. hVICs from passages 2–5 were used for the subsequent experiments in this study.

### 2.3 Osteogenic Induction

hVICs were seeded in plates and subsequently subjected to osteogenic induction using osteogenic medium (OM) supplemented with 10% fetal bovine serum, 1% penicillin‒streptomycin, 10 mmol/L beta-glycerophosphate (MedChemExpress, USA), 50 μg/mL ascorbic acid (MedChemExpress, USA), and 10 nmol/L dexamethasone (MedChemExpress, USA). After 21 days of culture, calcification was assessed by Alizarin Red S staining, and quantitative analysis was performed using ImageJ software.

### 2.4 Alizarin Red S Staining

After being washed twice with PBS, hVICs were fixed with 4% paraformaldehyde for 10 minutes. The cells were then rinsed twice with distilled water, followed by incubation with 0.2% Alizarin Red S solution (Servicebio, China) for 30 minutes in the dark. Excess dye was removed, and the cells were gently rinsed with distilled water to eliminate any residual stain. The calcified nodules were visualized and imaged under a microscope (EVOS M7000; Invitrogen, USA). Quantitative analysis was performed using ImageJ software to calculate the area of Alizarin Red S-positive staining.

### 2.5 Alkaline Phosphatase (ALP) Staining

The cells were subsequently washed twice with PBS and fixed with 4% paraformaldehyde for 10 minutes. After two additional washes with PBS, the cells were incubated with a staining solution containing BCIP, NBT, and alkaline phosphatase buffer (Beyotime, China) in the dark for 30 minutes. The staining solution was then removed, and the cells were washed twice with PBS. The stained cells were observed and imaged under a microscope. Quantitative analysis was performed using ImageJ software to calculate the area of ALP-positive staining.

### 2.6 Measurement of Calcium Content

Cultured cells were lysed with 0.1 M HCl and incubated at 4 °C overnight. The calcium concentration was quantified using a commercial calcium assay kit (Beyotime, China) based on the o-cresolphthalein complexone method. The calcium content was normalized to the total cellular protein amount and expressed as micrograms of calcium per milligram of protein.

### 2.7 Quantitative Real-Time PCR (qRT-PCR)

Total RNA was extracted from human aortic valve tissues using a commercial kit according to the manufacturer’s instructions. The quantity and quality of the RNA were determined, and equal amounts of RNA were reverse-transcribed into cDNA. Gene-specific primers were designed based on the cDNA sequences. qRT-PCR was performed using SYBR Green qPCR Mix (Sparkjade, China), with GAPDH serving as the internal control. Each reaction mixture contained SuperMix, nuclease-free water, specific primers, and template cDNA. The relative gene expression levels were calculated using the 2^-ΔΔCt^ method. The sequences of primers for the target genes are listed in Supplemental Table S2.

### 2.8 Western blot Analysis

Proteins were extracted from cells using RIPA lysis buffer (Sangon Biotech, China), followed by sonication and centrifugation at 4 °C to collect the supernatant. The protein concentration was quantified, and equal amounts of samples were denatured in 5× SDS loading buffer (Beyotime, China), separated by SDS-PAGE, and transferred onto PVDF membranes (Epizyme, China). After blocking with 5% non-fat milk in TBST for 1 h at room temperature, the membranes were incubated with primary antibodies overnight at 4 °C. Following three washes with TBST, the membranes were incubated with horseradish peroxidase-conjugated anti-mouse (1:5000) or anti-rabbit (1:10,000) secondary antibodies for 1 h at room temperature. The protein bands were visualized using an enhanced chemiluminescence (ECL) substrate (Epizyme, China) and imaged with a chemiluminescence detection system (Bio-Rad, USA). The band intensity was quantified using ImageJ software. The primary antibodies used in this study are listed in Supplemental Table S3.

### 2.9 CAVD Mouse Model

All animal experiments in this study were approved by the Animal Ethics Committee of Shandong First Medical University Affiliated Provincial Hospital (Approval No. SD NSFC 2025-400) and were conducted in accordance with the National Institutes of Health Guide for the Care and Use of Laboratory Animals. Male *ApoE^-/-^* mice were purchased from GemPharmatech (China). Mice were randomly divided into three groups (n=6 per group): one was fed a standard rodent chow diet, and the other two groups were fed a western diet for 24 weeks. One of the western diet groups received intraperitoneal injections of Y15 (30 mg/kg/day), while the other group received intraperitoneal injections of an equal volume of saline (see details in Major Resources Table). All the mice were housed in a specific pathogen-free, temperature-controlled environment under a 12-hour light/dark cycle with free access to food and water. Euthanasia was performed via intraperitoneal injection of a lethal dose of pentobarbital sodium (200 mg/kg). Prior to tissue collection, the mice were perfused with 4% paraformaldehyde via the left ventricle. Hearts were then dissected, embedded, and submitted to Bioss Biotech Co., Ltd., for sectioning.

### 2.10 Echocardiography

The mice were anesthetized via inhalation of 2.5% isoflurane, and echocardiographic parameters were acquired using a Vevo 2100 ultrasound system (VisualSonics Inc., Canada). Pulsed-wave Doppler was applied in the parasternal long-axis view to measure the peak aortic jet velocity, and the transvalvular pressure gradient was calculated. Measurements were averaged over at least three consecutive cardiac cycles. Images were obtained from the high right parasternal long-axis view, with the transducer angle maintained appropriately throughout the acquisition.

### 2.11 H&E Staining

Briefly, the paraffin sections were deparaffinized in xylene and rehydrated through a graded ethanol series. The sections were then stained in haematoxylin solution (Servicebio, China) for 5–10 minutes, rinsed with tap water, and subjected to differentiation and bluing. The sections were subsequently counterstained in 0.5% eosin solution for 1–3 minutes. After dehydration through a graded ethanol series, the sections were cleared in xylene twice (5 minutes each) and mounted with neutral resin. The stained tissue sections were observed and imaged under a microscope.

### 2.12 Masson’s Trichrome Staining

Masson’s Trichrome Staining of the paraffin sections was performed using a commercial kit (Solarbio, China) according to the manufacturer’s instructions. Briefly, the sections were stained with Weigert’s iron haematoxylin solution for 5 minutes. After being rinsed with distilled water, the sections were treated with acid differentiation solution for 10 seconds, followed by Masson’s blue solution for 5 minutes. The sections were subsequently stained with Ponceau S/Fuchsin solution for 5 minutes, rinsed briefly with a weak acid working solution, and sequentially treated with phosphomolybdic acid solution and aniline blue solution, with rinses in the weak acid working solution between steps. Finally, the sections were dehydrated through a graded ethanol series, cleared in xylene, mounted with neutral resin, and placed under a microscope for image acquisition.

### 2.13 Von Kossa Staining

Von Kossa staining of the paraffin sections was performed using a commercial kit (Servicebio, China) according to the manufacturer’s instructions. Briefly, the slides were placed in a humidified chamber, and the tissue was covered with Von Kossa staining solution. The slides were exposed to ultraviolet light for 4 hours. Following incubation, the excess staining solution was removed by washing with ultrapure water. Finally, the sections were dehydrated in absolute ethanol, cleared in xylene, mounted with neutral resin, and imaged under a microscope.

### 2.14 Co-immunoprecipitation (Co-IP)

Co-immunoprecipitation was performed as previously described^28^. Briefly, cell lysates were incubated with antibody-bound protein A+G beads (Beyotime, China) at room temperature on a shaker or rotary mixer for 2 hours, after which the beads were magnetically separated and then washed three times with lysis buffer containing protease inhibitors. The immunoprecipitated complexes were eluted, denatured, and subjected to SDS-PAGE analysis.

### 2.15 Immunofluorescence

After the tissue sections were deparaffinized in xylene and rehydrated through a graded ethanol series, antigen retrieval was performed in a microwave oven at 97 °C for 30 minutes. Endogenous peroxidase activity was blocked with 3% hydrogen peroxide for 10 minutes. The sections were then incubated with wheat germ agglutinin (WGA) (Servicebio, China) for 20 minutes, followed by incubation with primary antibody for 1 hour. After being washed with TBST, the sections were incubated with a fluorescent-conjugated secondary antibody for 30 minutes. Nuclei were counterstained with 4’,6-diamidino-2-phenylindole (DAPI).

After two washes with PBS, the cells were fixed with 4% paraformaldehyde for 10 minutes. The cells were then permeabilized and blocked. The membranes were subsequently incubated with primary antibody at 4 °C overnight, followed by incubation with a fluorescent secondary antibody for 1 hour at room temperature in the dark. Nuclei were stained with DAPI. Stained samples were observed and imaged under a fluorescence microscope (DMi8; Leica, Germany). Quantitative analysis of the fluorescence signal area was performed using ImageJ software. The primary antibodies used in this study are listed in Supplemental Table S3.

### 2.16 Preparation of Decellularized Extracellular Matrix (dECM)

When the cells reached 80–90% confluence, they were treated with one of the following solutions for 5 minutes: 1) 0.25% SDS, 2) 0.5% SDS, 3) 0.25% SDS + 0.5% Triton X-100, or 4) 0.5% SDS + 0.5% Triton X-100. After treatment, the cell layers were thoroughly washed with PBS. The resulting dECM was stored at 4 °C for up to one week for subsequent use.

### 2.17 Scanning Electron Microscopy

dECM samples were processed for scanning electron microscopy. The samples were fixed with 2.5% glutaraldehyde, dehydrated through a graded ethanol series, substituted with isoamyl acetate and subjected to critical point drying. After drying, the samples were sputter-coated with gold and observed under a scanning electron microscope (Regulus8100, Hitachi, Japan) at an accelerating voltage of 5–20 kV to examine their surface morphology and microstructure.

### 2.18 Proteomics

Quantitative proteomic analysis was performed using liquid chromatography-tandem mass spectrometry (LC-MS/MS). Briefly, aortic valve tissue samples were ground in liquid nitrogen and lysed to extract total protein. The protein concentration was determined using a BCA assay. Equal amounts of protein were subjected to reduction, alkylation, and tryptic digestion. The resulting peptides were separated by chromatography using a NanoElute system and analysed on a timsTOF Pro mass spectrometer operating in parallel accumulation–serial fragmentation (PASEF) mode. The raw MS data were processed by database searching and bioinformatics analysis to identify differentially expressed proteins and infer their biological functions.

### 2.19 RNA Sequencing

Transcriptome sequencing was performed on six samples, comprising the VASP-OE group and the Control group, with three biological replicates per group. High-throughput sequencing was conducted on Illumina, and the raw image data were converted into raw reads in FASTQ format via base calling. Gene expression levels were quantified using StringTie software, and normalized using both Fragments Per Kilobase of transcript per Million mapped reads (FPKM) and Transcripts Per Million (TPM) methods. Differential expression analysis between groups (Control vs. VASP-OE) was performed using the DESeq2 package. Genes meeting the threshold of a P-value < 0.05 and an absolute log_2_-fold change |log_2_FC| ≥ 1 were identified as differentially expressed genes (DEGs). To explore the biological functions and signalling pathways associated with the DEGs, Functional Enrichment analyses were conducted. GO Functional Enrichment: DEGs were mapped to the Gene Ontology (GO) database to identify significantly enriched terms (P ≤0.05) across three categories: biological process (BP), cellular component (CC), and molecular function (MF). KEGG Pathway Enrichment: Hypergeometric tests were performed using the clusterProfiler package to identify biochemical metabolic pathways or signal transduction pathways significantly enriched with DEGs. Furthermore, a protein-protein interaction (PPI) network was constructed by importing the DEGs into the STRING database to identify core regulatory hubs.

### 2.20 Statistical Analysis

For comparisons between two groups, Student’s t-test or the Mann-Whitney U test was applied. For comparisons among three or more groups, one-way analysis of variance (ANOVA) followed by Bonferroni’s multiple comparisons test was used. Quantitative data are presented as the mean ± standard deviation and represent at least three independent experiments. A P-value ≤ 0.05 was considered to indicate statistical significance. All the statistical calculations were performed using GraphPad software.

## 3. Results

### 3.1 VASP is significantly upregulated in calcified aortic valves

To identify proteins highly expressed in calcified aortic valve tissues, we performed proteomic profiling on calcified and non-calcified aortic valve specimens (3 vs. 3) and identified 93 significantly upregulated proteins (Fig. 1A). We subsequently intersected this list with RNA-sequencing data from the GEO database (GSE235995). To ensure a baseline expression level, parameters were set as base mean ≥ 500, fold change ≥ 2, and p value < 0.05. The intersection analysis revealed seven overlapping genes: VASP, ITGB2, CTSZ, CTSB, FN1, FHL2, and FERMT3 (Fig. 1A). Further validation by RT-qPCR revealed that FN1, FHL2, and VASP were significantly upregulated in calcified aortic valve tissues, with VASP being the most significantly upregulated (p≤0.001) (Fig. 1B). In addition, VASP expression in human aortic valve tissues was assessed through Western blotting and immunofluorescence. The findings revealed a significant increase in VASP expression in calcified aortic valve tissues (Fig. 1C, D). Furthermore, upregulation of VASP expression was also observed in the aortic valve tissues of western diet-fed *ApoE^-/-^* mice (Fig. 1E). Next, Next, we examined the expression changes of VASP in VICs at different time points during osteogenic induction *in vitro*. The results showed that the osteogenic markers RUNX2 and ALP were both significantly upregulated in a time-dependent manner, reaching the highest level on day 5 (Fig. 1F). Meanwhile, VASP was also significantly upregulated with a similar time-dependent pattern, but its expression essentially reached the highest level on day 3 of induction (Fig. 1F). Collectively, these data demonstrate that VASP is highly expressed in calcified aortic valve tissues and in osteogenic-induced VICs *in vitro*, suggesting a potentially important role in CAVD.

**Figure 1.**
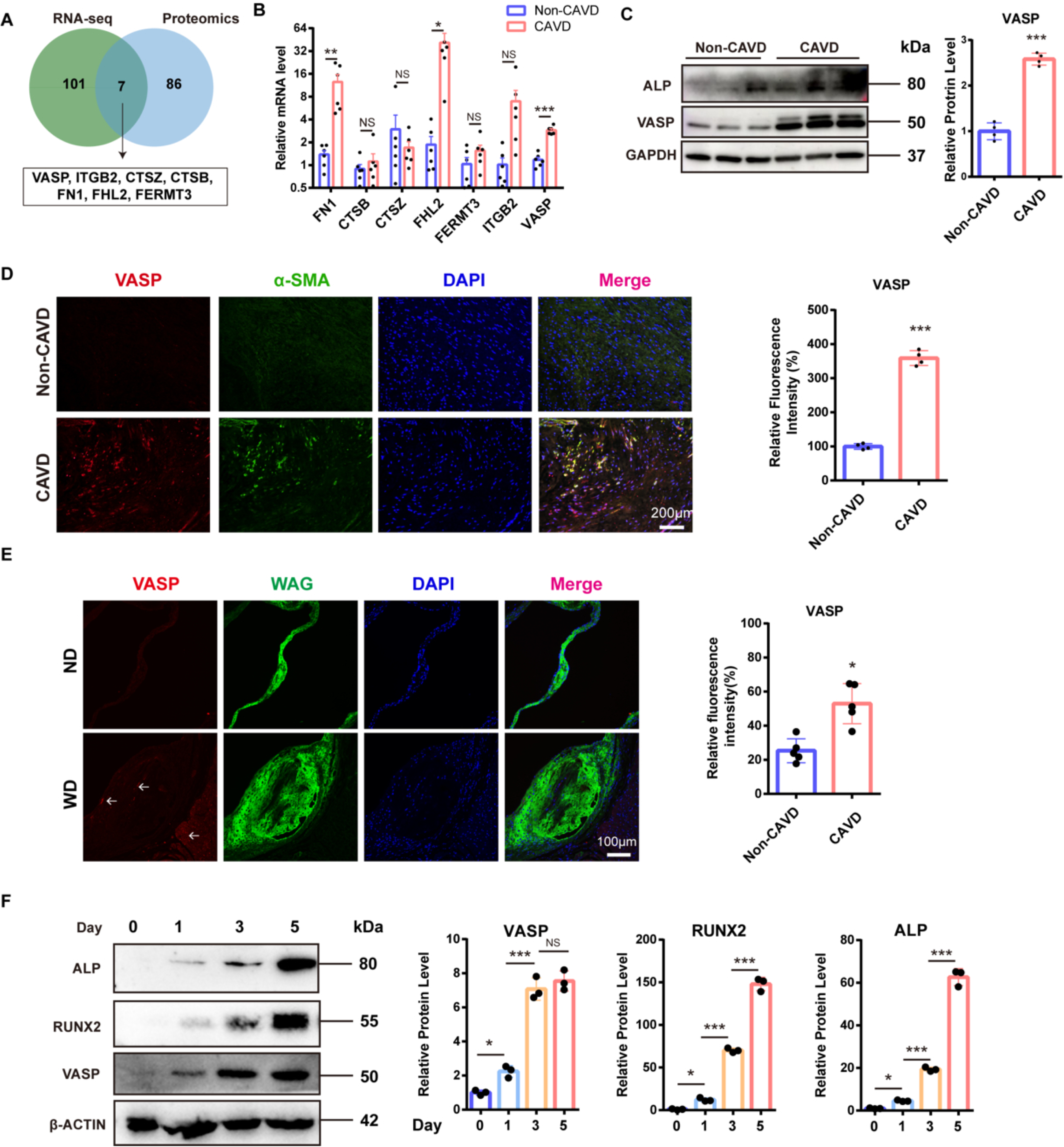
VASP is highly expressed in calcified aortic valve tissues. **(A)** Venn diagram comparing the upregulated differentially expressed genes (DEGs) from proteomic analysis and the GEO dataset (GSE235995). **(B)** Validation of the upregulated genes by RT-qPCR (n=6). **(C, D)** VASP is highly expressed in aortic valve tissues from CAVD patients (n=4). **(E)** VASP is upregulated in the aortic valve tissues of western diet-fed *ApoE^⁻/⁻^* mice (n=5; scale bar: 100 μm). **(F)** RUNX2, ALP and VASP were all significantly upregulated in a time-dependent manner (n=3). The values are the means ± SDs. *P* values were calculated using an unpaired two-tailed Student’s t test. NS: not significant, *p<0.05, **p<0.01, ***p<0.001; ND: normal diet; WD: Western diet

### 3.2 VASP regulates the osteogenic differentiation of VICs in vitro

Osteogenic differentiation of VICs is considered a central event in CAVD. To investigate the role of VASP in CAVD, we knocked down VASP in VICs and cultured the cells in osteogenic induction medium (OM). Alizarin Red S staining revealed that knockdown of VASP reduced the area of calcified nodules in VICs (Fig. 2A, B). ALP staining revealed that VASP knockdown decreased the ALP-positive area (Fig. 2C, D). Measurement of calcium content further confirmed that VASP knockdown inhibited calcium deposition (Fig. 2E). Western blot analysis of osteogenic markers revealed that VASP knockdown suppressed the expression of RUNX2 and ALP (Fig. 2F).

**Figure 2.**
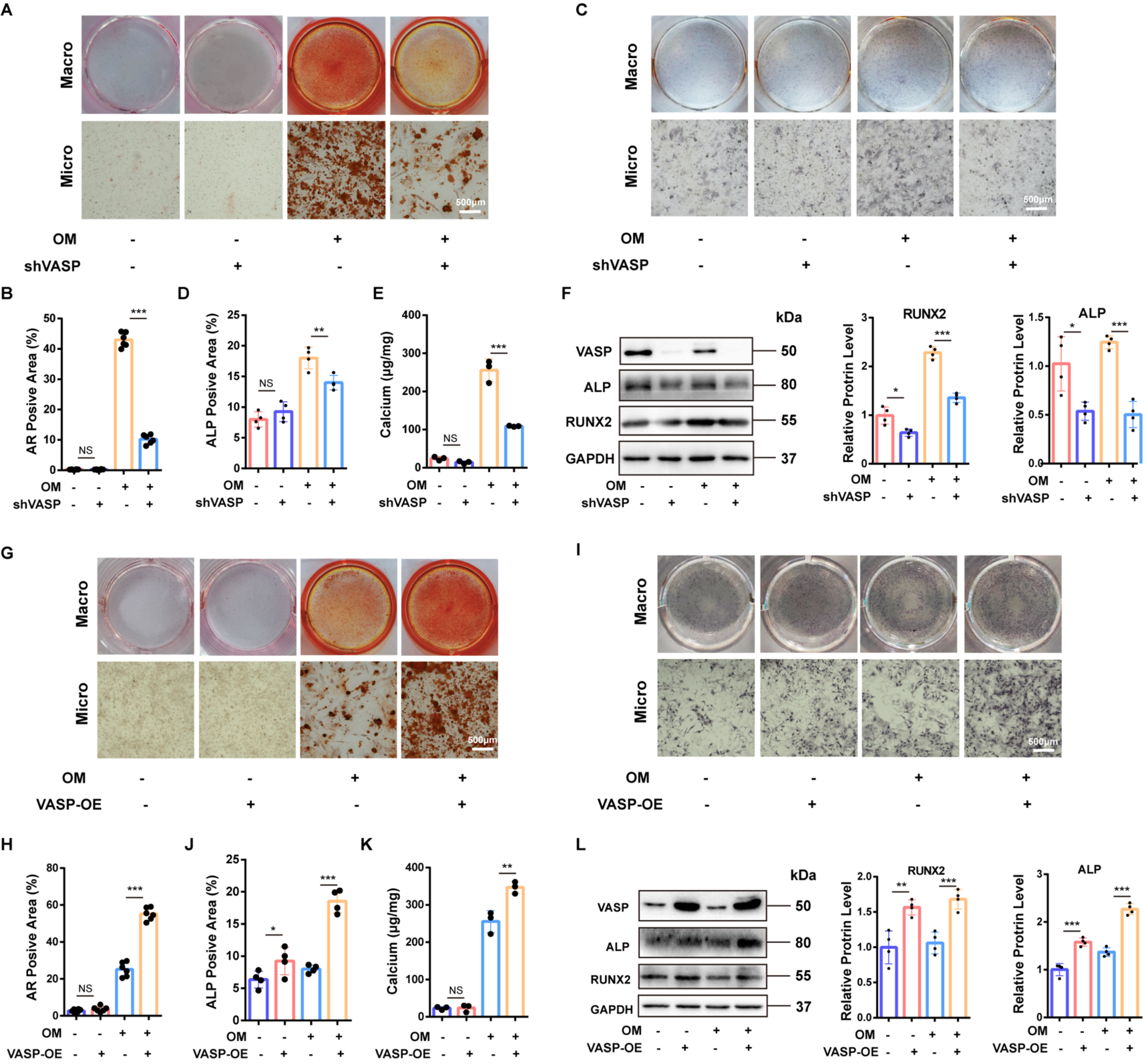
VASP positively regulates the osteogenic differentiation of VICs. (A, **B)** Knockdown of VASP reduces calcified nodule formation (n=6; scale bar: 500 µm). **(C, D)** Knockdown of VASP decreases the ALP-positive area (n=4; scale bar: 500 µm). **(E)** Knockdown of VASP reduces calcium content (n=3). **(F)** Knockdown of VASP downregulates the expression of RUNX2 and ALP (n=4). **(G, H)** Overexpression of VASP promotes calcified nodule formation (n=6; scale bar: 500 µm). **(I, J)** Overexpression of VASP increases the ALP-positive area (n=4; scale bar: 500 µm). **(K)** Overexpression of VASP increases the calcium concentration (n=3). **(L)** Overexpression of VASP upregulates the expression of RUNX2 and ALP (n=4). The values are the means ± SDs. *P* values were calculated using one-way ANOVA followed by the Bonferroni multiple comparisons test; NS: not significant; *p<0.05, **p<0.01, ***p<0.001

To further confirm the regulatory effect of VASP on the osteogenic differentiation of VICs, we overexpressed VASP in these cells. VASP overexpression increased the formation of calcified nodules (Fig. 2G, H) and the ALP-positive area (Fig. 2I, J), increased the calcium concentration (Fig. 2K) and upregulated the expression of the osteogenic markers RUNX2 and ALP (Fig. 2L) in VICs. These results reveal that VASP positively regulates the osteogenic differentiation of VICs in vitro.

### 3.3 VASP promotes osteogenic differentiation of VICs by activating the FA pathway

To explore the mechanism by which VASP promotes osteogenic differentiation of VICs, we performed RNA sequencing on VICs overexpressing VASP (3 vs. 3). Seventy-two genes were upregulated, and 58 were downregulated (Fig. 3A). Gene Ontology (GO) analysis revealed that VASP overexpression was associated with biological processes such as plasma membrane formation, heart development, myofibril formation, and positive regulation of osteoblast differentiation (Fig. 3B). KEGG pathway analysis revealed that the differentially expressed genes were predominantly enriched in the focal adhesion (FA) signalling pathway (Fig. 3C). We then examined the expression of p-FAK, a key molecule in the FA pathway, and found that VASP overexpression increased p-FAK levels (Fig. 3D, E).

**Figure 3.**
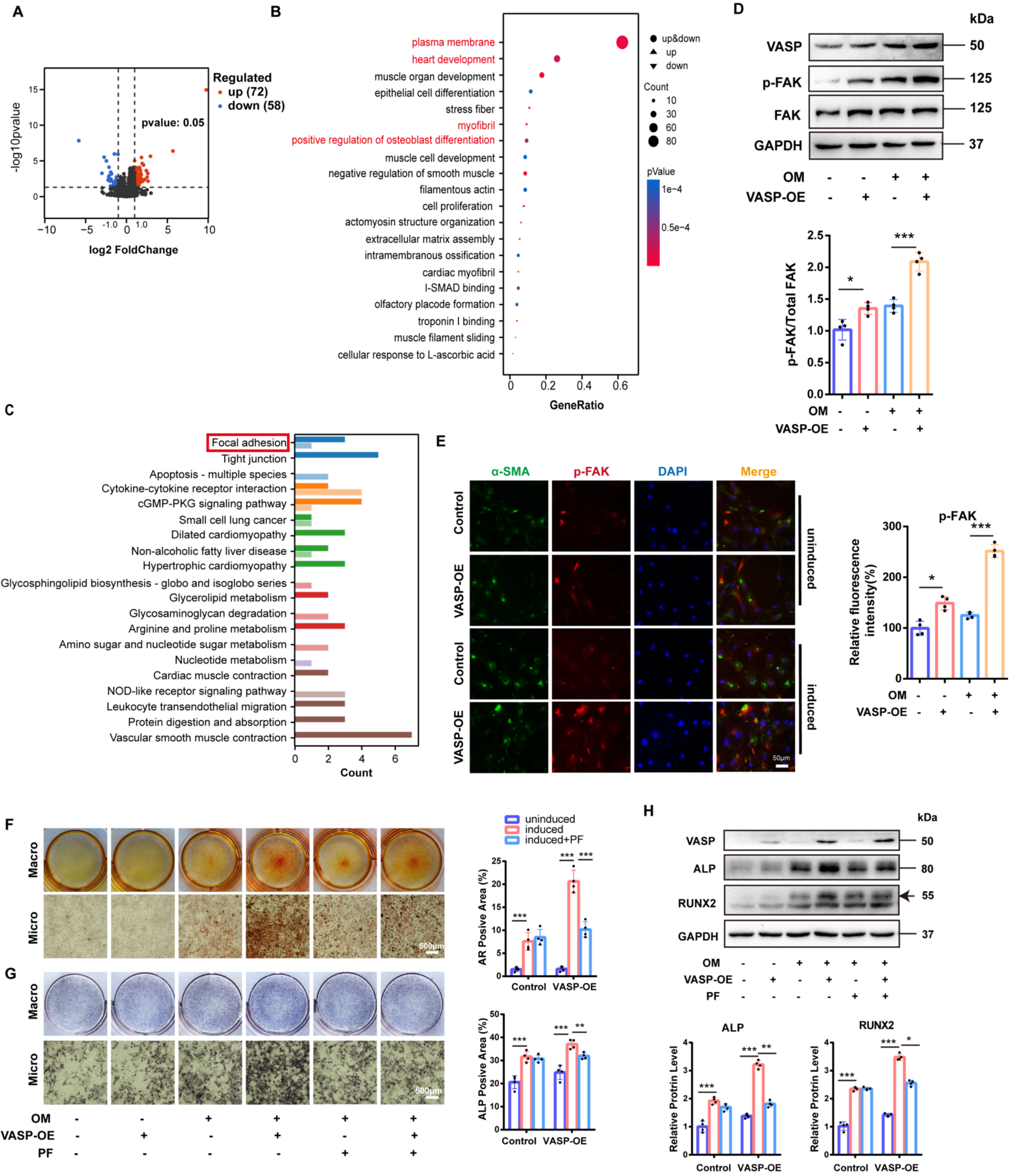
The FA pathway mediates the pro-osteogenic effect of VASP. **(A)** Volcano plot of the differentially expressed genes. **(B)** GO enrichment analysis. **(C)** KEGG enrichment analysis. **(D, E)** Overexpression of VASP increases FAK phosphorylation (n=4; scale bar: 50 µm). **(F)** PF-573228 significantly inhibits calcified nodule formation (n=4; scale bar: 500 µm). **(G)** PF-573228 significantly inhibits ALP activity (n=4; scale bar: 500 µm). **(H)** PF-573228 significantly inhibits the expression of ALP and RUNX2 (n=4). The values are the means ± SDs. *P* values were calculated using one-way ANOVA followed by the Bonferroni multiple comparisons test; *p<0.05, **p<0.01, ***p<0.001

To further determine whether the pro-osteogenic effect of VASP is mediated by the FA pathway, we supplemented the OM with the FA pathway inhibitor PF-573228 during osteogenic induction. CCK-8 assay showed that PF573228 at concentrations below 500 nM did not affect the viability of VICs (Fig. S1). Therefore, we used 100 nM PF573228 for subsequent experiments. PF-573228 suppressed the VASP overexpression–induced increase in calcified nodule formation (Fig. 3F) and ALP activity (Fig. 3G) and inhibited the upregulation of osteogenic markers (Fig. 3H). These results indicate that the FA pathway mediates the pro-osteogenic effect of VASP on VICs.

### 3.4 Inhibition of the FA pathway attenuates the calcification phenotype in a CAVD mouse model

A CAVD model was established by feeding male *ApoE^-/-^* mice a Western diet for 24 weeks. Immunofluorescence staining revealed increased p-FAK expression in the aortic valve tissues of Western diet–fed mice, which was significantly reduced by treatment with the FA pathway inhibitor Y15 (Fig. S2), indicating that the FA pathway is activated in the CAVD mouse model and that Y15 effectively inhibits this activation. Echocardiography revealed that Western diet feeding significantly increased the peak aortic jet velocity and transvalvular pressure gradient, effects that were markedly suppressed by Y15 treatment (Fig. 4A, B). Histological staining revealed significant leaflet thickening, increased collagen deposition, and enhanced calcification in Western diet-fed *ApoE^-/-^* mice. Y15 treatment effectively reversed these changes (Fig. 4C–E). In addition, the upregulation of the osteogenic marker BMP2 was reversed by Y15 (Fig. 4F). Furthermore, biochemical analysis showed that Y15 treatment did not alter the levels of triglycerides, total cholesterol, high-density lipoprotein cholesterol, or low-density lipoprotein cholesterol in Western diet-fed *ApoE^⁻/⁻^* mice, but significantly reduced blood glucose levels (Fig. S3A-E). These results reveal that inhibition of the FA pathway attenuates the calcification phenotype in the CAVD mouse model.

**Figure 4.**
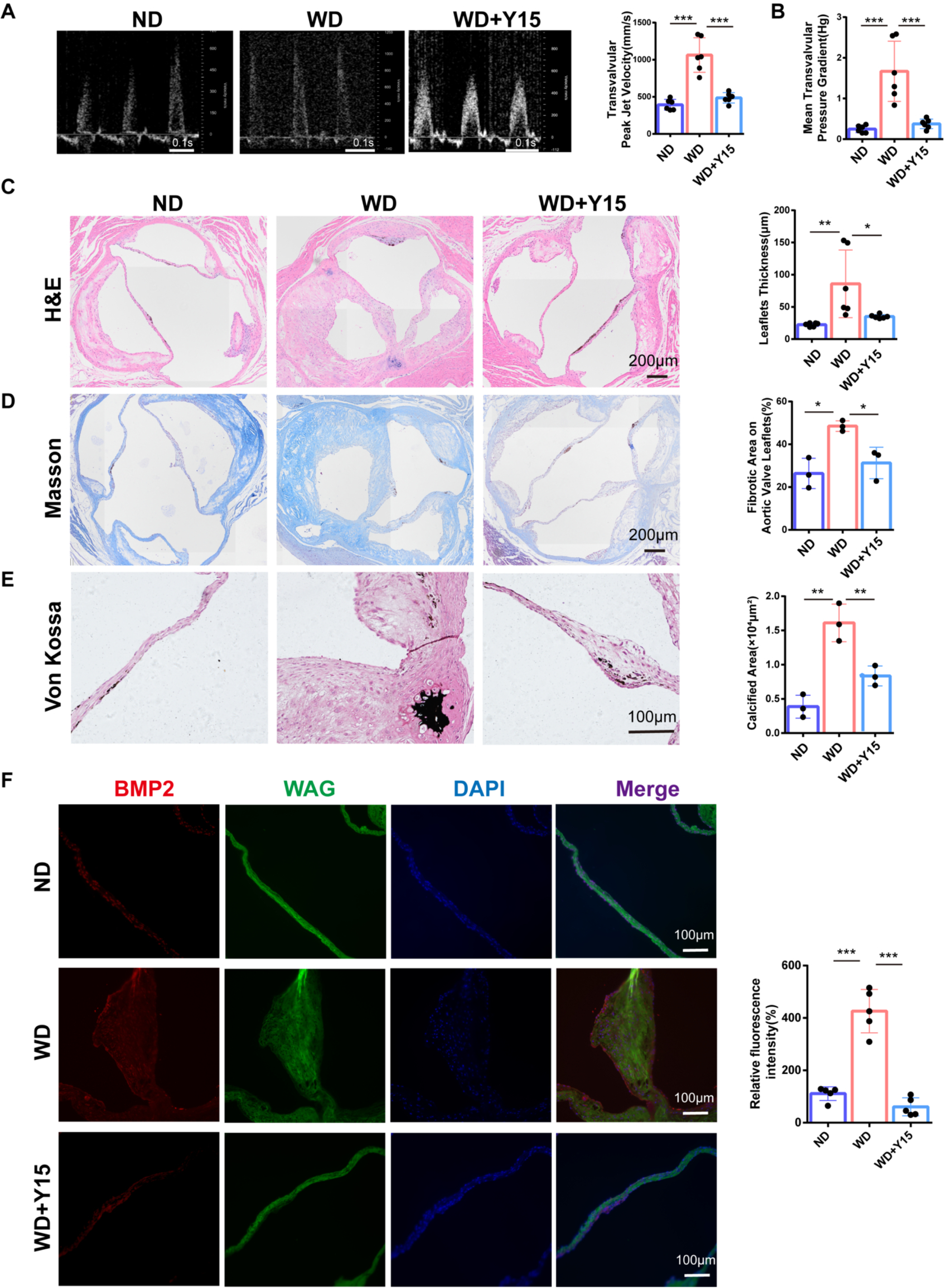
Supplementation with an FA pathway inhibitor attenuates the calcification phenotype in a CAVD mouse model. **(A)** Y15 treatment significantly reduces the peak aortic jet velocity in Western diet–fed mice (n=6). **(B)** Y15 treatment significantly reduces the mean transvalvular pressure gradient in Western diet–fed mice (n=6). **(C)** Y15 treatment significantly reduces aortic valve leaflet thickness in Western diet–fed mice (n=6; scale bar: 200 µm). **(D)** Y15 treatment significantly reduces the fibrotic area in the aortic valve tissues of Western diet–fed mice (n=3; scale bar: 200 µm). **(E)** Y15 treatment significantly reduces the calcified area in the aortic valve tissues of Western diet–fed mice (n=3; scale bar: 100 µm). **(F)** Y15 treatment significantly reduces BMP2 expression in the aortic valve tissues of Western diet–fed mice (n=5; scale bar: 100 µm). The values are the means ± SDs. *P* values were calculated using one-way ANOVA followed by the Bonferroni multiple comparisons test; *p<0.05, **p<0.01, ***p<0.001

### 3.5 FBLIM1 participates in the pro-osteogenic process by interacting with VASP

To explore the molecular mechanism through which VASP activates the FA pathway, we searched for VASP-interacting proteins in the STRING database, which revealed a predicted interaction between VASP and FBLIM1 (Fig. 5A). Correlation analysis revealed a significant positive relationship between the expression levels of VASP and FBLIM1 (Fig.5B,C). This interaction was further confirmed by coimmunoprecipitation (Fig. 5D) and molecular docking simulation, with the latter suggesting the formation of hydrogen bonds between the two proteins (Fig. 5E). FBLIM1 contains two disordered regions and three LIM zinc-binding domains (Fig. 5F). To identify the specific domain of FBLIM1 involved in binding to VASP, we cotransfected 293T cells with Flag-tagged VASP and various His-tagged full-length or truncated FBLIM1 constructs. Co-IP assays demonstrated that the interaction was abolished upon deletion of the third LIM zinc-binding domain (Fig. 5G), indicating that this domain mediates the binding to VASP.

**Figure 5.**
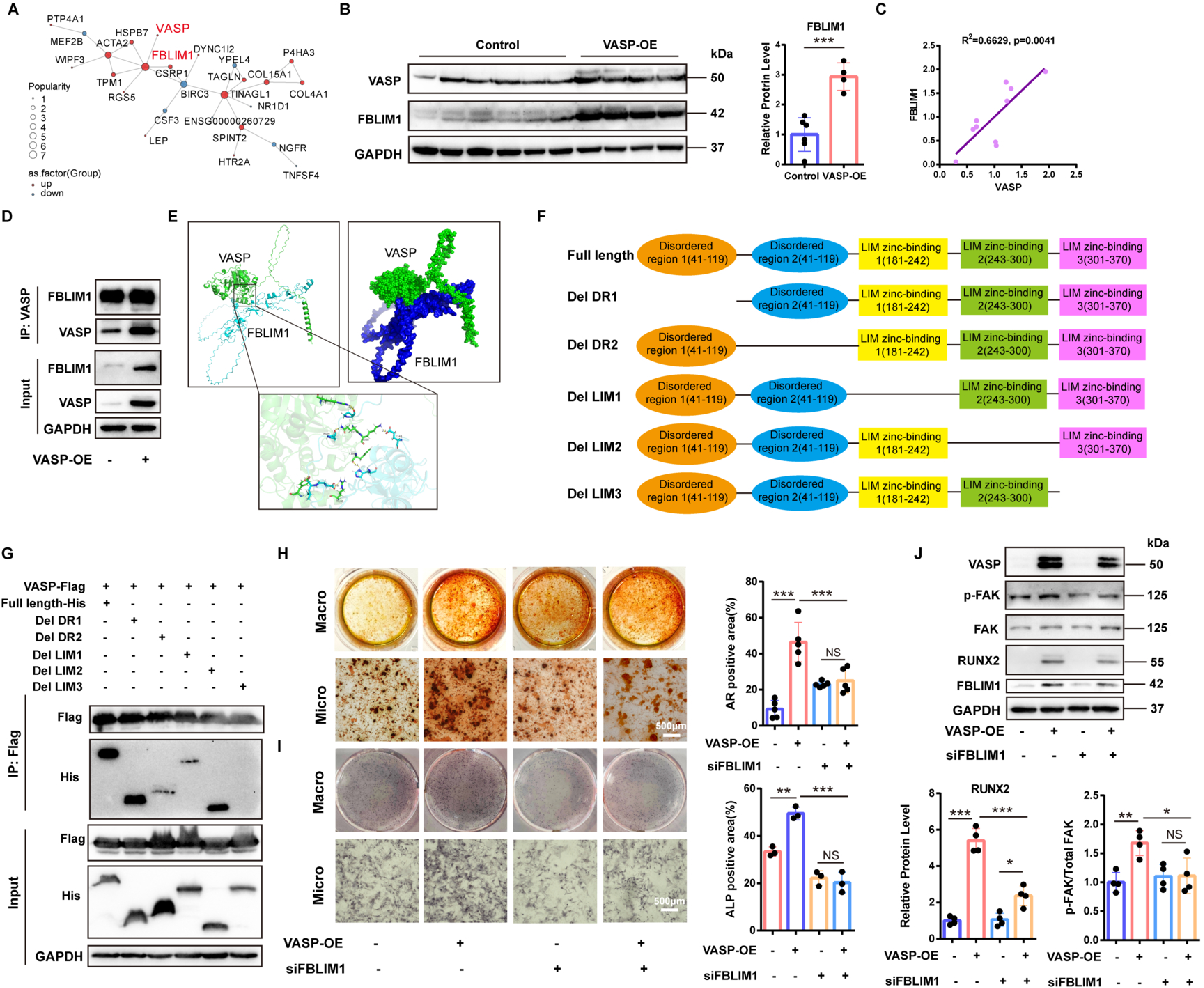
FBLIM1 interacts with VASP and activates the FA pathway. **(A)** Analysis of the STRING database reveals an interaction between VASP and FBLIM1. **(B)** Overexpression of VASP upregulates the expression of FBLIM1 (Control: n=6, VASP-OE: n=4). **(C)** FBLIM1 expression is positively correlated with VASP expression (two-tailed Pearson correlation analysis, n=10). **(D)** VASP and FBLIM1 interact with each other (Co-IP). **(E)** VASP and FBLIM1 form hydrogen bonds (molecular docking). **(F)** Schematic representation of the full-length and truncated domains of FBLIM1. **(G)** VASP binds to the third LIM zinc-binding domain of FBLIM1 (Co-IP). **(H)** Knockdown of FBLIM1 abolishes the VASP overexpression–induced increase in calcified nodule formation (n=5; scale bar: 500 µm). **(I)** Knockdown of FBLIM1 abolishes the VASP overexpression–induced increase in the ALP-positive area (n=3; scale bar: 500 µm). **(J)** Knockdown of FBLIM1 attenuates the VASP overexpression–induced upregulation of RUNX2 expression and inhibits p-FAK expression (n=4). The values are the means ± SDs. *P* values were calculated using one-way ANOVA followed by the Bonferroni multiple comparisons test; NS: not significant; *p<0.05, **p<0.01, ***p<0.001

To determine whether FBLIM1 is involved in the pro-osteogenic effect of VASP, we knocked down FBLIM1 in VICs overexpressing VASP. FBLIM1 knockdown reduced calcium deposition and calcified nodule formation (Fig. 5H), decreased the ALP-positive area (Fig. 5I), and downregulated RUNX2 expression (Fig. 5J), demonstrating that FBLIM1 participates in VASP-mediated osteogenic differentiation. Furthermore, FBLIM1 knockdown inhibited p-FAK expression (Fig. 5J), indicating that FBLIM1 mediates the activation of the FA pathway.

### 3.6 The VASP/FA pathway promotes osteogenic differentiation of VICs by enhancing ELN secretion

Focal adhesions are dynamic structures that link cells to the extracellular matrix (ECM), facilitating bidirectional signalling via integrin receptors. CAVD progression is accompanied by increased ECM production and deposition. To investigate whether VASP regulates ECM deposition by activating the FA pathway, we analysed gene expression changes. A gene heatmap revealed that VASP overexpression upregulated the expression of the ECM components COL4A1 and ELN (Fig. 6A), which was validated by qPCR (Fig. 6B). Western blot analysis further confirmed the increase in ELN expression upon VASP overexpression, and this effect was reversed by the FA inhibitor PF-573228 (Fig. 6C). In contrast, COL4A1 levels did not significantly change (Fig. 6C), suggesting that the VASP/FA pathway primarily regulates ELN expression. Consistent with these findings, ELN expression was upregulated in the aortic valve tissues of the CAVD mouse model, and this upregulation was suppressed by FA pathway inhibition (Fig. 6D). These results indicate that ELN is positively regulated by the VASP/FA axis.

**Figure 6.**
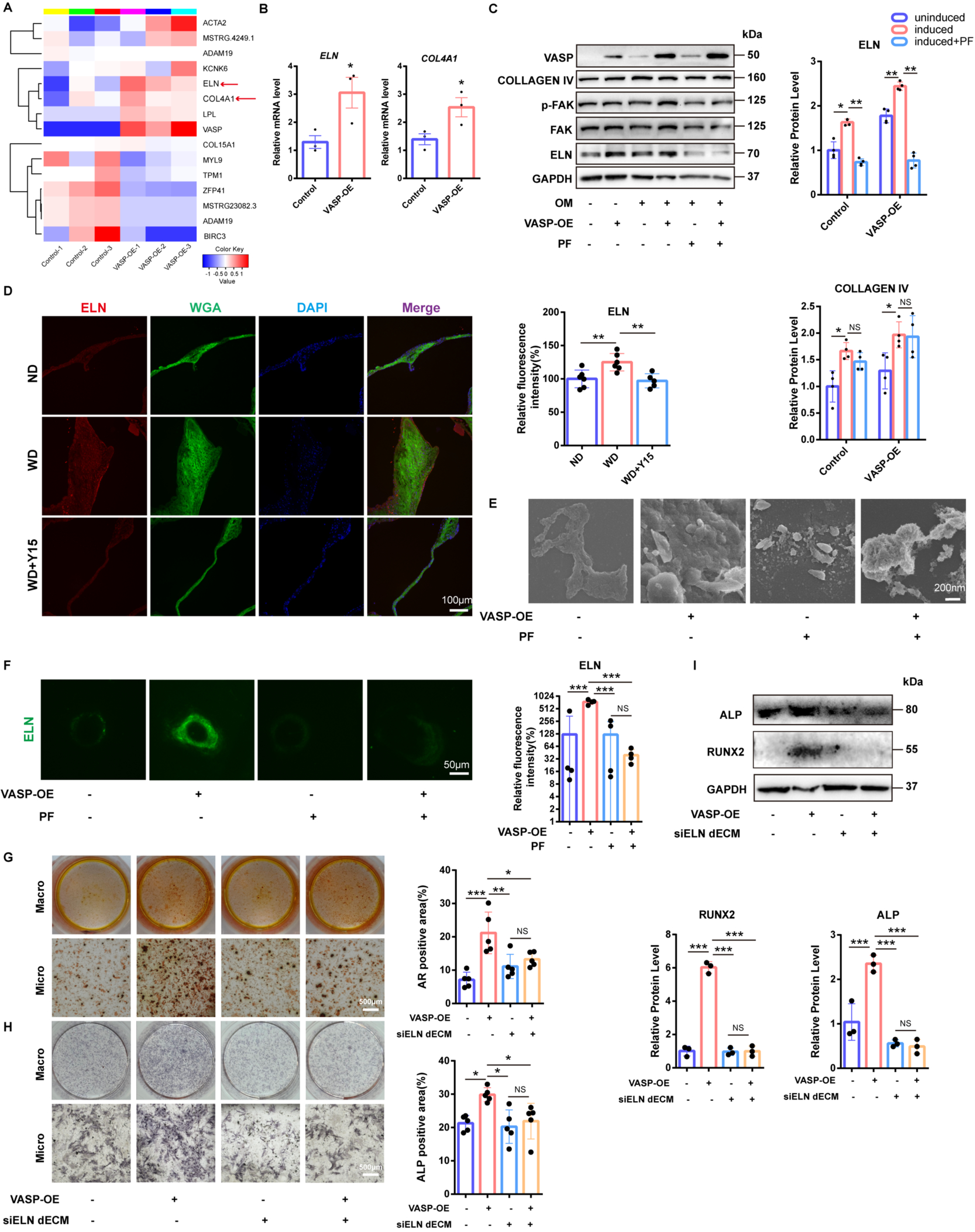
The VASP/FA Pathway Promotes the Osteogenic Differentiation of VICs by Enhancing ELN Secretion. (A, **B)** Overexpression of VASP upregulates the expression of COL4A1 and ELN (n=3). **(C)** PF-573228 abolishes the VASP overexpression–induced upregulation of ELN expression (n=4). **(D)** Y15 abolishes the upregulation of ELN expression in the aortic valves of Western diet–fed mice (n=6; scale bar: 100 µm). **(E)** PF-573228 treatment reduces the VASP overexpression–induced increase in ECM deposition (scanning electron microscopy). **(F)** PF-573228 treatment reduces the VASP overexpression–induced increase in ELN secretion (n=4; scale bar: 50 µm). **(G)** Knockdown of ELN abolishes the VASP overexpression–induced increase in calcified nodule formation (n=5; scale bar: 500 µm). **(H)** Knockdown of ELN abolishes the VASP overexpression–induced increase in the ALP-positive area (n=5; scale bar: 500 µm). **(I)** Knockdown of ELN abolishes the VASP overexpression–induced upregulation of RUNX2 and ALP expression (n=3). The values are the means ± SDs. *P* values were calculated using one-way ANOVA followed by the Bonferroni multiple comparisons test; NS: not significant; *p<0.05, **p<0.01, ***p<0.001

To determine whether ELN expression affects the osteogenic differentiation of VICs, we decellularized VICs to obtain cell-derived ECM. Treatment with 0.25% SDS plus 0.5% Triton X-100 yielded optimal decellularization, as evidenced by better preservation of ECM components (Fig. S4A, B) and more complete DNA removal (Fig. S4C). Scanning electron microscopy of dECM from VICs overexpressing VASP, with or without PF-573228 treatment, revealed that VASP overexpression increased ECM secretion, which was reduced by PF-573228 treatment (Fig. 6E). Immunofluorescence staining of the dECM confirmed that PF-573228 treatment significantly decreased ELN deposition (Fig. 6F).

To investigate the functional impact of ELN, we seeded fresh VICs onto dECM derived from VICs that overexpressed VASP with or without ELN knockdown, followed by osteogenic induction. VICs grown on dECM from VASP-overexpressing cells exhibited increased calcified nodule formation (Fig. 6G), ALP activity (Fig. 6H), and osteogenic marker expression (Fig. 6I). In contrast, these pro-osteogenic effects were reversed when ELN was knocked down in the donor VICs (Fig. 6G-I). Collectively, these results reveal that the VASP/FA pathway promotes VIC osteogenic differentiation by enhancing the secretion of ELN.

## 4. Discussion

The pathological progression of calcific aortic valve disease (CAVD) has been established as an actively regulated process primarily driven by the aberrant activation of valvular interstitial cells (VICs). However, the early mechanotransduction mechanisms and the initiating factors of extracellular matrix (ECM) remodelling remain incompletely understood. In this study, for the first time, we reveal the central driving role of vasodilator-stimulated phosphoprotein (VASP) in CAVD pathogenesis. Our key findings reveal that VASP is significantly upregulated in diseased valve tissues and, through direct interaction with FBLIM1, aberrantly activates the focal adhesion (FA) signalling pathway. This activation subsequently promotes the excessive secretion of elastin (ELN), ultimately leading to the osteogenic differentiation of VICs and valvular calcification. In vitro experiments confirmed that VASP knockdown inhibited the osteogenic differentiation of VICs, whereas its overexpression markedly promoted a calcification phenotype. More importantly, FA pathway inhibitors effectively reversed VASP-mediated VIC osteogenesis and attenuated the calcification phenotype in a CAVD mouse model. Collectively, these results reveal the pivotal role of the VASP-FBLIM1-FA-ELN axis in CAVD progression, providing a novel perspective for understanding its pathophysiology.

VASP, as a crucial actin-binding protein, plays multiple roles in cytoskeletal remodelling and signal transduction^29^. This study explores how VASP regulates the osteogenic differentiation of VICs through its molecular interaction network. We identified a direct physical interaction between VASP and FBLIM1, with FBLIM1 binding to VASP via its third LIM zinc-finger domain. FBLIM1, a known focal adhesion-associated protein, is involved in cytoskeletal organization and signalling^30^. The binding of VASP to FBLIM1 may alter the structure and function of focal adhesions, thereby activating the downstream FA pathway, particularly the phosphorylation of FAK. The activated FA pathway further promotes ECM remodelling, especially the oversecretion of ELN. ELN is a major component of the aortic valve, and its abnormal accumulation and cross-linking may provide a scaffold for calcium salt deposition, thereby facilitating the development of CAVD. Thus, the VASP-FBLIM1-FA-ELN cascade constitutes a sophisticated molecular regulatory network in CAVD pathogenesis.

The findings of this study are highly consistent with the growing literature emphasizing the role of the mechanical microenvironment in cardiovascular diseases^31–36^. For instance, Cao et al. reported that increased matrix stiffness is an independent risk factor for the induction of the osteogenic differentiation of VICs^34^. Our conclusions not only corroborate this point but also further identify the VASP-FBLIM1 axis as a critical molecular switch mediating this mechanical response. However, in contrast to the traditional view that collagen types I and III are the primary contributors to the fibrotic stage, this study unexpectedly reveals that the abnormal, substantial deposition of collagen type IV and ELN plays a dominant role in the early remodelling mediated by VASP. This discrepancy may stem from our focus on a relatively early stage of CAVD progression (the transition from aortic valve sclerosis to calcification), during which the turnover of basement membrane-associated components is higher, and the abnormal synthesis and subsequent fragmentation/degradation of ELN may serve as initial nucleation sites for calcification. This discovery not only addresses a gap in the literature but also suggests that different ECM proteins may play dynamically shifting roles across various stages of CAVD.

Despite these important advances, this study has several limitations. First, its primary focus is on the VASP-FBLIM1-FA-ELN axis. CAVD is a complex, multifactorial disease involving multiple pathways^37–41^, and VASP may influence CAVD through other, yet undiscovered, mechanisms. Future research could employ more comprehensive omics technologies, such as single-cell sequencing, to elucidate the heterogeneous roles of VASP in different VIC subpopulations. Second, While the western diet-fed ApoE^-/-^mouse model effectively recapitulates many features of human fibro-calcific valve remodelling, it does not perfectly mirror the hemodynamics or severe calcific stenosis seen in elderly clinical populations. Due to the systemic nature of our pharmacological intervention and the current lack of a valve-specific, conditionally targeted VASP^-/-^mouse model, we cannot entirely exclude systemic off-target actions or secondary metabolic influences on the observed valvular phenotype. Future studies utilizing lineage-tracing systems or leaflet-specific gene manipulation will be essential to definitively cement the localized therapeutic potential of targeting this axis. And validation in larger human cohorts and prospective clinical studies is needed to assess the therapeutic potential of targeting VASP or the FA pathway. Furthermore, from a mechanistic standpoint, while we successfully mapped the direct binding of VASP to the third LIM zinc-binding domain of FBLIM1, the precise structural basis and downstream structural dynamics of this complex under variable mechanical strains warrant further crystalline or cryo-EM characterization. Finally, our focus on focal adhesion kinase (FAK) phosphorylation as a primary signalling readout represents only one facet of a highly complex mechanotransduction network. The broader roles of other focal adhesion components, such as paxillin, vinculin, and interconnected cytoskeletal forces, were not exhaustively explored in this study.

In summary, this study is the first to reveal the key mechanism through which VASP promotes the osteogenic differentiation of VICs and CAVD progression through interaction with FBLIM1 to activate the FA pathway, thereby driving excessive ELN secretion. These findings not only strengthen our understanding of CAVD pathophysiology but also provide promising targets for the development of novel diagnostic and therapeutic strategies.

## Authors’ contributions

Z.J.W., and X.L.L. conceived of and designed the study. X.L.L., Q.Y.X., and K.X. performed the experiments *in vitro* and analyzed data. Q.Z., N.Z., and W.L.Z. provided technical guidance. K.X., and P.L.S. established an animal model. R.L., and Z.S.L. collected clinical samples.

## Acknowledgements

We appreciate all patients all who donated their aortic valves. We also thank Prof. Dong Wang for providing clinical samples.

## Supplemental Material

Figures S1–S4

Tables S1-S3

Major Resources Table

